# Systematic functional characterization of the intellectual disability-associated SWI/SNF complex reveals distinct roles for the BAP and PBAP complexes in post-mitotic memory forming neurons of the *Drosophila* mushroom body

**DOI:** 10.1101/408500

**Authors:** Melissa C. Chubak, Max H. Stone, Nicholas Raun, Shelby L. Rice, Mohammed Sarikahya, Spencer G. Jones, Taylor A. Lyons, Taryn E. Jakub, Roslyn L.M. Mainland, Maria J. Knip, Tara N. Edwards, Jamie M. Kramer

## Abstract

Technology has led to rapid progress in the identification of genes involved in neurodevelopmental disorders like intellectual disability (ID), but our functional understanding of the causative genes is lagging. Here, we show that the SWI/SNF chromatin remodeling complex is one of the most overrepresented cellular components disrupted in ID. We systematically investigated the role of individual subunits of this large protein complex in post-mitotic memory forming neurons of the *Drosophila* mushroom body (MB). Using this approach, we have identified novel differential roles for the two prominent conformations of the *Drosophila* SWI/SNF complex, known as BAP and PBAP. The PBAP conformation is required post-mitotically for remodeling of the MB γ neurons during morphogenesis and is essential for both short and long-term memory. In contrast, the BAP conformation appears to preferentially effect long-term memory and is associated with γ neuron survival. Our results suggest that different subunits of the SWI/SNF complex may influence learning and memory through diverse and distinct roles in regulating structural plasticity, survival, and functionality of post-mitotic neurons. This study provides novel insight into the neuronal function of individual SWI/SNF subunits and will serve as a basis for understanding SWI/SNF-mediated gene regulatory mechanisms in post-mitotic neurons.

## Introduction

Intellectual disability (ID) is a neurodevelopmental disorder characterized by early onset limitations in cognitive function and adaptive behaviour that affects 1-3% of the global population. Technological advances in DNA sequencing have led to rapid progress in understanding the genetic etiology of ID, and as a result, there are currently almost 1000 known primary ID genes (sysid.cmbi.umcn.nl/)^1^. For most of these genes, we have no knowledge of their role in the nervous system. Therefore, functional analysis of known ID genes is the next frontier in advancing our understanding of neurodevelopmental disorders.

About two thirds of known ID genes follow a recessive or X-linked inheritance pattern. Historically, such inheritance patterns have made it possible to identify disease genes using family pedigrees combined with genomic methodologies, like homozygosity mapping. However, recent studies suggest that recessive and X-linked inheritance patterns are not representative of the majority of ID cases^2-7^. In fact, most people with ID have a dominant genetic variant that is acquired *de novo*. Dominant *de novo* copy number variations and single nucleotide variants are estimated to account for 60% of severe ID cases, compared to only 2% of severe ID cases that are caused by rare inherited forms^2^. Several large-scale studies on cohorts comprised of patients with variable clinical presentation paint a similar picture, with a prominent role for dominant ID genes (DIGs)^4,5,7,8^.

Recently, all known ID genes have been documented in a hand curated publicly accessible database called sysID (sysid.cmbi.umcn.nl)^1^. There are 339 dominant DIGs documented in this database (update Dec 2017). Here, we show that DIGs are highly cohesive, suggesting that they may be involved in common pathways or biological processes. We find that DIGs are enriched for genes encoding proteins associated with chromatin regulation and identify the SWI/SNF ATP-dependent chromatin remodelling complex as the most enriched DIG-associated cellular component. Currently, mutations in 9 of the 29 genes that encode subunits of the human SWI/SNF complex have been found in patients with ID^7–11^.

The SWI/SNF complex was originally identified in yeast and is highly conserved^9^. Each conformation of the SWI/SNF complex contains 10-15 protein subunits, including a single ATPase that utilizes energy from ATP to alter nucleosome positioning, making chromatin either more or less accessible for interactions with transcription factors. Recently, the SWI/SNF complex has been shown to increase chromatin accessibility at cell type specific enhancers in human cancer cell lines and mouse embryonic fibroblasts^10-12^. The chromatin remodeling action of the SWI/SNF complex is essential in maintaining global epigenetic programs required for cell type specification, differentiation, and neuronal stem cell proliferation during mouse brain development^13-18^. Double knockout of the paralogous SWI/SNF components BAF155 and BAF170 in mouse leads to proteasomal degradation of the entire complex, providing a unique system to investigate the consequences of complete loss of SWI/SNF function in neurons^14^. In this scenario, it was shown that SWI/SNF is critical in neural progenitors for the specification of different brain structures^14,19^.

These studies highlight the essential nature of SWI/SNF mediated gene regulation in neuronal differentiation, however the mechanisms that are disrupted in ID are still not known. ID-causing SWI/SNF mutations are heterozygous and have been proposed to cause haploinsufficiency, dominant negative, or gain of function effects depending on the nature of the mutation and the specific SWI/SNF subunit involved^20-23^. It is likely that partial SWI/SNF function is maintained in individuals with these mutations. Therefore, despite the clear importance of SWI/SNF subunits in cell differentiation and tissue specification, human SWI/SNF related disorders may in part result from more subtle defects in gene regulation in post-mitotic neurons. In mouse, the Baf53b SWI/SNF subunit is only incorporated into the complex in differentiated post-mitotic neurons. Deletion of this neuron specific subunit does not affect complex assembly or neuron survival, but is required for activity dependent dendritic outgrowth, long-term memory, and for regulation of specific genes^24-26^. Like Baf53b, other SWI/SNF components are also known to mediate specific gene regulatory mechanisms without interfering with the overall integrity of the complex^27,28^. However, the role of most SWI/SNF complex components in post-mitotic neurons remains unexplored.

Here, we have systematically characterized the role of individual SWI/SNF subunits through targeted RNAi knockdown in post-mitotic neurons of the *Drosophila* mushroom body (MB), a critical brain structure for learning and memory^29,30^. The SWI/SNF complex is highly conserved in *Drosophila* and has two distinct configurations known as BAP (brahma associated proteins) and PBAP (polybromo BAP) ^31^. BAP and PBAP contain an overlapping set of core subunits as well as the complex-specific subunits; osa for BAP, and polybromo, Bap170, and e(y)3, for PBAP (**Figure 2B**) ^32,33^. We have identified and characterized unique requirements for individual SWI/SNF components in neuron remodeling, cell survival, and memory, in post-mitotic memory forming neurons of the *Drosophila* MB.

**Figure 1:**
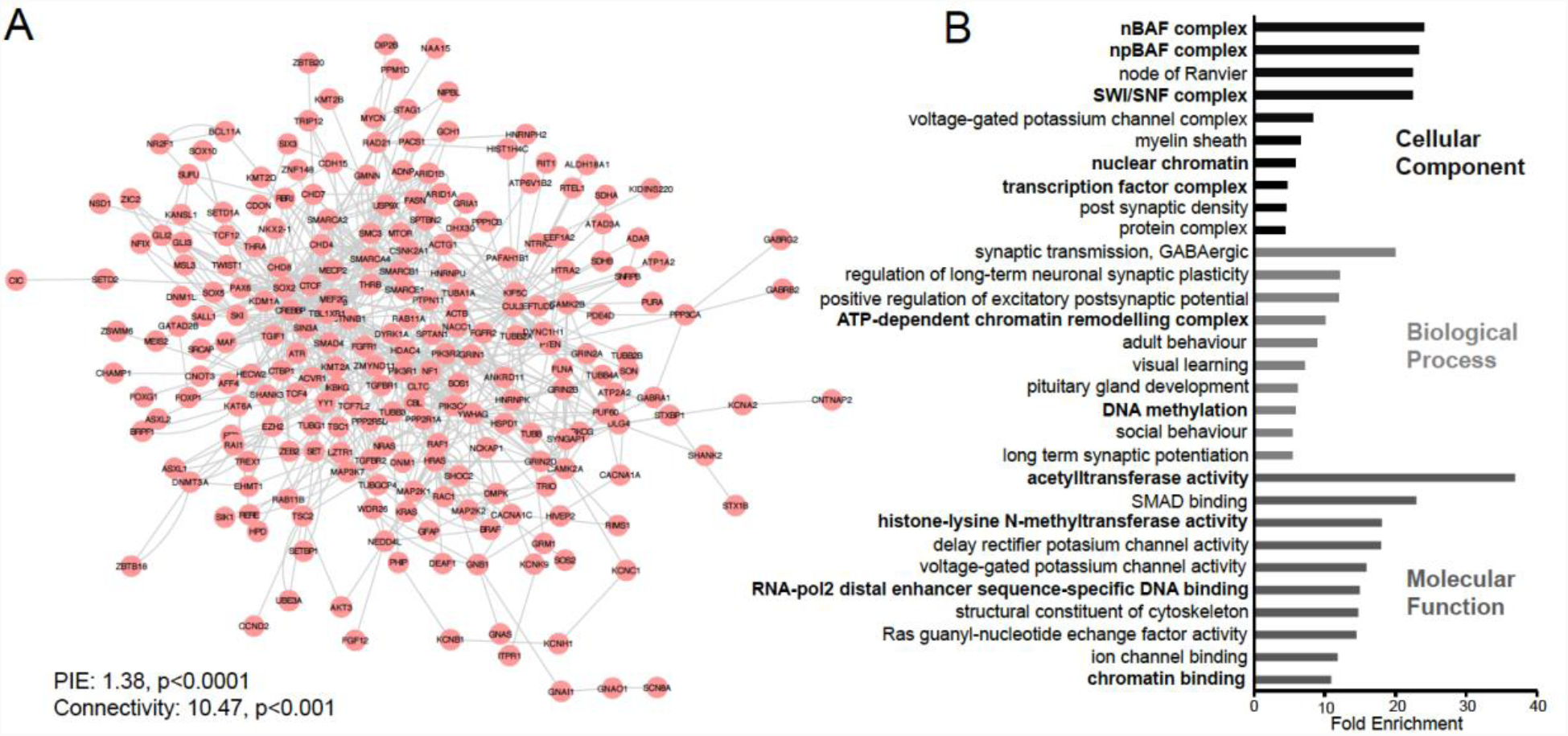
Dominant ID genes are highly cohesive and enriched for the SWI/SNF chromatin remodeling complex. (**A**) Protein interaction network of 339 dominant primary ID genes (DIGs) obtained from sysID (sysid.cmbi.umcn.nl). 235 DIGs form a single network based on annotated protein-protein interactions in BioGrid small scale studies and the Human Protein Reference Database (HPRD). DIGs have significantly more interactions and connectivity than expected by random chance (PIE algorithm). (**B**) Gene ontology enrichment analysis for 339 DIGs. Top 10 enriched terms with a Bonferroni corrected p-value < 0.05 for each GO category are shown. Terms related to gene and chromatin regulation are shown in bold.

**Figure 2:**
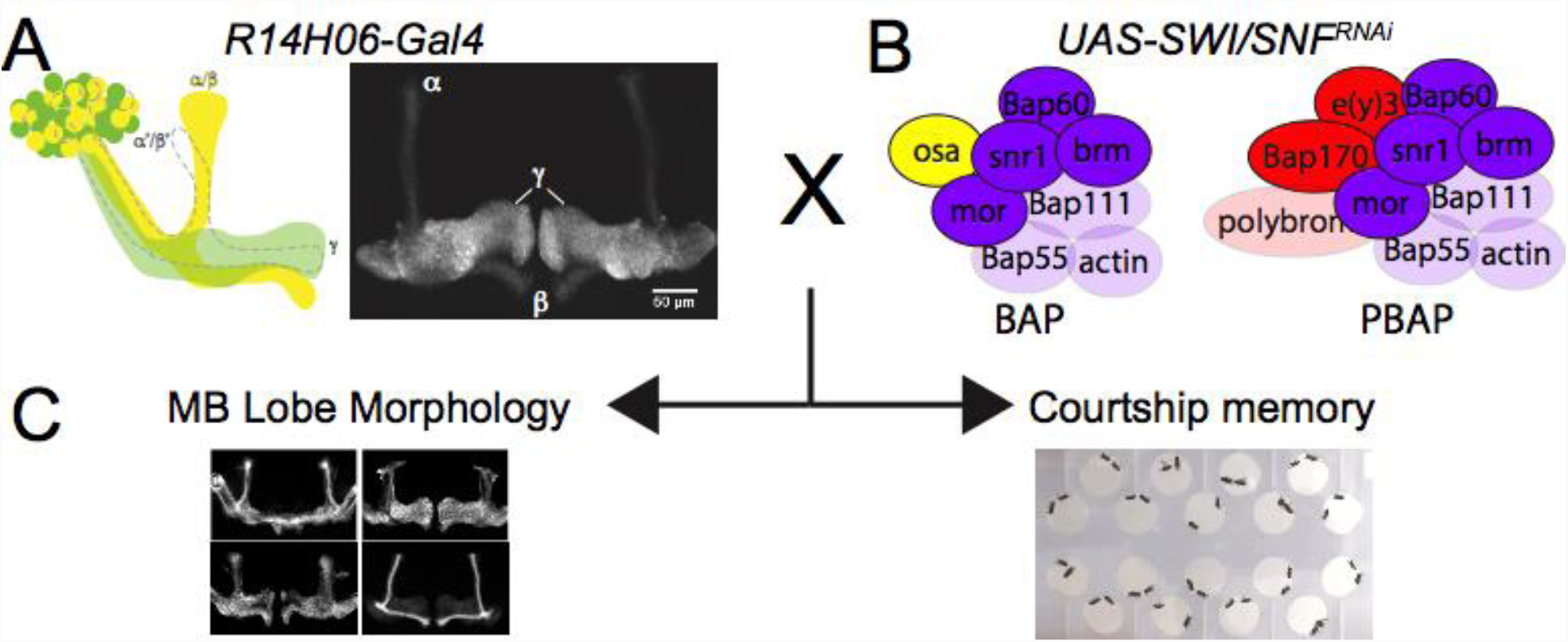
Schematic diagram indicating the experimental strategy. The mushroom body specific (MB) Gal4 driver *R14H06-Gal4* (**A**) was used to express *UAS-RNAi* lines targeting 7 different components of the *Drosophila* SWI/SNF complex (**B**). SWI/SNF knockdown flies and controls were examined for defects in MB morphology, and courtship memory. (**A**) Schematic diagram and confocal projection showing the expression domain of *R14H06-Gal4*. A full brain confocal stack is available in **Video S1. (B)** Schematic representation of the BAP and PBAP conformations of the SWI/SNF complex. Blue – core subunits, yellow – BAP specific subunits, red – PBAP specific subunits. Subunits with validated RNAi lines used in this study are indicated with solid colour. Other subunits are indicated by transparent colour.

## Results

### The SWI/SNF complex is the most enriched cellular component among DIGs

Considering that most ID cases are caused by dominant *de novo* mutations, we investigated whether known DIGs possess any common functionality. Using sysID^1^ we retrieved a list of 339 known primary DIGs. Using data from BioGrid^34^ and the Human Protein Reference Database^35^, we asked whether DIGs are involved in annotated protein-protein interactions (PPIs) with each other. Strikingly, 235 of the 339 genes form a single PPI network (**Fig. 1A**). Statistical analysis using the protein interaction enrichment (PIE) algorithm^36^ shows that the number of PPIs between DIGs is almost 40% more than would be expected with a random set of proteins with an equal number of known PPIs (PIE value = 1.38, p < 0.0001). Also, the connectivity of the DIG PPI network is nearly 11-fold higher than expected (p<0.001). This implies that DIGs are highly cohesive and may operate in similar biological processes or pathways. To further investigate the potential overlapping function of DIGs, we performed gene ontology enrichment analysis. Enriched terms related primarily to neuronal components and functions (**Fig. 1B, non-bold terms**), or chromatin regulation (**Fig. 1B, bold terms**). The most enriched gene ontology terms for cellular components were related specifically to the SWI/SNF chromatin remodeling complex and one of the most enriched terms for biological processes was “ATP-dependent chromatin remodeling complex”. This demonstrates that disruption of chromatin regulation is a major factor in the etiology of ID, and identifies the SWI/SNF complex as the most overrepresented protein complex associated with ID. The SWI/SNF complex is known to be essential for differentiation and cell type specification in neuronal progenitors (for review see^13^). However, individuals with SWI/SNF mutations do not appear to have major defects in differentiation or tissue specification, suggesting that SWI/SNF likely has important roles in post-mitotic neurons. Aside from the Baf53b subunit, which is exclusively expressed in post-mitotic neurons in mice^15^, little is known about the role of individual SWI/SNF subunits in differentiated neurons. Therefore, we set out to systematically investigate the function of SWI/SNF components in post-mitotic memory forming neurons in *Drosophila*.

### Establishment of tools for investigation of SWI/SNF components in *Drosophila* memory forming neurons

Since null mutations in most SWI/SNF complex components cause embryonic lethality, we chose to specifically target memory forming neurons of the *Drosophila* MB using the UAS/Gal4 binary expression system in combination with flies containing Gal4 inducible UAS-RNAi transgenes (see methods). MB specificity was achieved using the Gal4 driver *R14H06-Gal4*, which expresses Gal4 specifically in post-mitotic cells contributing to the γ and α/β lobes of the MB (**Fig. 2A, Supplementary video 1**)^37^. We aimed to test two RNAi constructs targeting different regions of the mRNA for 10 established *Drosophila* SWI/SNF subunits: brm, Bap60, snr1, Bap111, mor, osa, e(y)3, Bap170, Bap55, and polybromo (**Figure 2B**). Although two RNAi lines were available for polybromo, one of them (VDRC stock number 108618) did not survive well under normal culture conditions. For the remaining genes, we initially tested RNAi efficiency by measuring percent survival of adult progeny when RNAi transgenes were expressed ubiquitously using *Actin-Gal4* (**Table S1**). Of 19 RNAi lines tested, 15 caused near complete lethality (survival < 5%). Two RNAi lines were excluded (*UAS-Bap55*^31708^ and *UAS-brm*^34520^), because they did not decrease the rate of survival compared to controls. An additional 2 RNAi lines showed an intermediate percent survival that was still statistically less than controls; 53% for *UAS-Snr1-RNAi*^12644^, and 17% for *UAS-Bap60-RNAi*^33954^ (**Table S1**). For most RNAi transgenes, we performed additional validation of RNAi efficiency by qPCR upon ubiquitous knockdown in whole larvae. We observed a decrease in mRNA using *UAS-brm-RNAi*^37720^ and *UAS-brm-RNAi*^31712^ (**Table S1**), each of which cause lethality in combination with the ubiquitous *Actin-Gal4* driver line^38^. In contrast, the *UAS-brm*^34520^ RNAi line, which caused no reduction in survival in combination with *Actin-Gal4*, also caused no reduction in mRNA levels compared to controls (**Table S1**). This suggests that the lethality assay is a good proxy towards judging the efficiency of the RNAi lines used in this study. All other RNAi lines that were tested by qPCR did show a significant reduction in mRNA levels compared to controls with the exception of *Bap111*. Despite inducing lethality upon expression with Act-Gal4, *UAS-Bap111*^35242^ showed increased mRNA levels by qPCR, while *UAS-Bap111*^26218^ showed no change in mRNA levels. These results were confirmed at the protein level by Western blot (**Figure S1**), and therefore, Bap111 was excluded from further analysis. Here we focus our investigation on the 7 SWI/SNF subunits for which we had two validated RNAi lines. This included 4 core SWI/SNF components (brm, Bap60, Snr1, and mor), the BAP-specific subunit osa, and the PBAP-specific subunits e(y)3 and Bap170 (**Figure 2B**). For each RNAi line we looked for defects in MB morphology and memory (**Figure 2C**).

### SWI/SNF components in the regulation of mushroom body morphogenesis

The *R14H06-Gal4* driver is expressed in postmitotic neurons of the MB γ and α/β lobes (**Figure 2A**). γ neurons begin to arise in the early larval stage while α/β neurons arise during pupal development^39^. It was possible that SWI/SNF knockdown might affect any post-mitotic processes effecting MB morphology, such as axonogenesis. Therefore, SWI/SNF knockdown MBs were examined for gross morphological defects using confocal microscopy on dissected whole mount adult brains. This revealed five distinct phenotypic classes that were observed at various frequencies across the 348 adult fly brains that were imaged as part of this experiment. These phenotypes included: 1) missing α and β lobes, 2) β-lobe fibers crossing the midline, 3) stunted γ-lobes 4) extra dorsal projections, and 5) faded γ-lobes. Phenotypes were qualitatively assessed for severity and compared statistically to relevant control genotypes (see methods). The first two phenotypic classes, missing α and β lobes, and β-lobe midline crossing, were observed across many knockdown genotypes and controls at a low frequency. The appearance of missing α and β lobes was very rare, occurring in 2.7% of control brains and 3.0% of knockdown brains. In addition, there was no significant difference between knockdown and control genotypes for any RNAi line (**Figure S2**). The β-lobe crossing phenotype was observed more frequently with 12.6% of control brains and 18.7% of knockdown brains showing a phenotype, usually of mild or moderate severity (**Figure S3**). However, there were no cases where two RNAi lines targeting the same gene caused a significant increase in the occurrence of this phenotype. Sporadic appearance of missing lobes and β-lobe crossing phenotypes were previously reported in a study looking at variation in MB morphology across different genetic backgrounds^40^. Therefore, taken together with our findings, it seems unlikely that these phenotypes are specifically related to SWI/SNF RNAi knockdown.

The next two observed phenotypic classes, stunted γ lobes and extra dorsal projections, appeared to coincide in certain knockdown genotypes at a high penetrance. RNAi lines targeting Bap60, Snr1 and e(y)3 resulted in a near complete penetrance of the extra dorsal projection phenotype, which was significantly greater than that seen in controls (**Figure 3A and 3B**). This phenotype was consistent between two RNAi lines for Bap60 and e(y)3, suggesting that it is not likely due to off-target RNAi effects. For Snr1, the phenotype was not consistent between two RNAi lines tested, with *UAS-Snr1*^12644^ causing no phenotype. This discrepancy is likely a result of the weaker knockdown efficiency observed with *UAS-Snr1*^12644^, which showed a 53% survival rate in our *Actin-Gal4* induced lethality assay, compared to 4% for *UAS-Snr1*^32372^ (**Table S1**). For Bap60 and Snr1 RNAi lines, the appearance of a severe extra dorsal projection phenotype corresponded with the presence of stunted γ lobes indicating that these two phenotypes may be mechanistically linked (**Figure S4**). Overall, these findings suggest that Bap60, Snr1, and e(y)3 may regulate specific aspects of MB morphogenesis.

**Figure 3:**
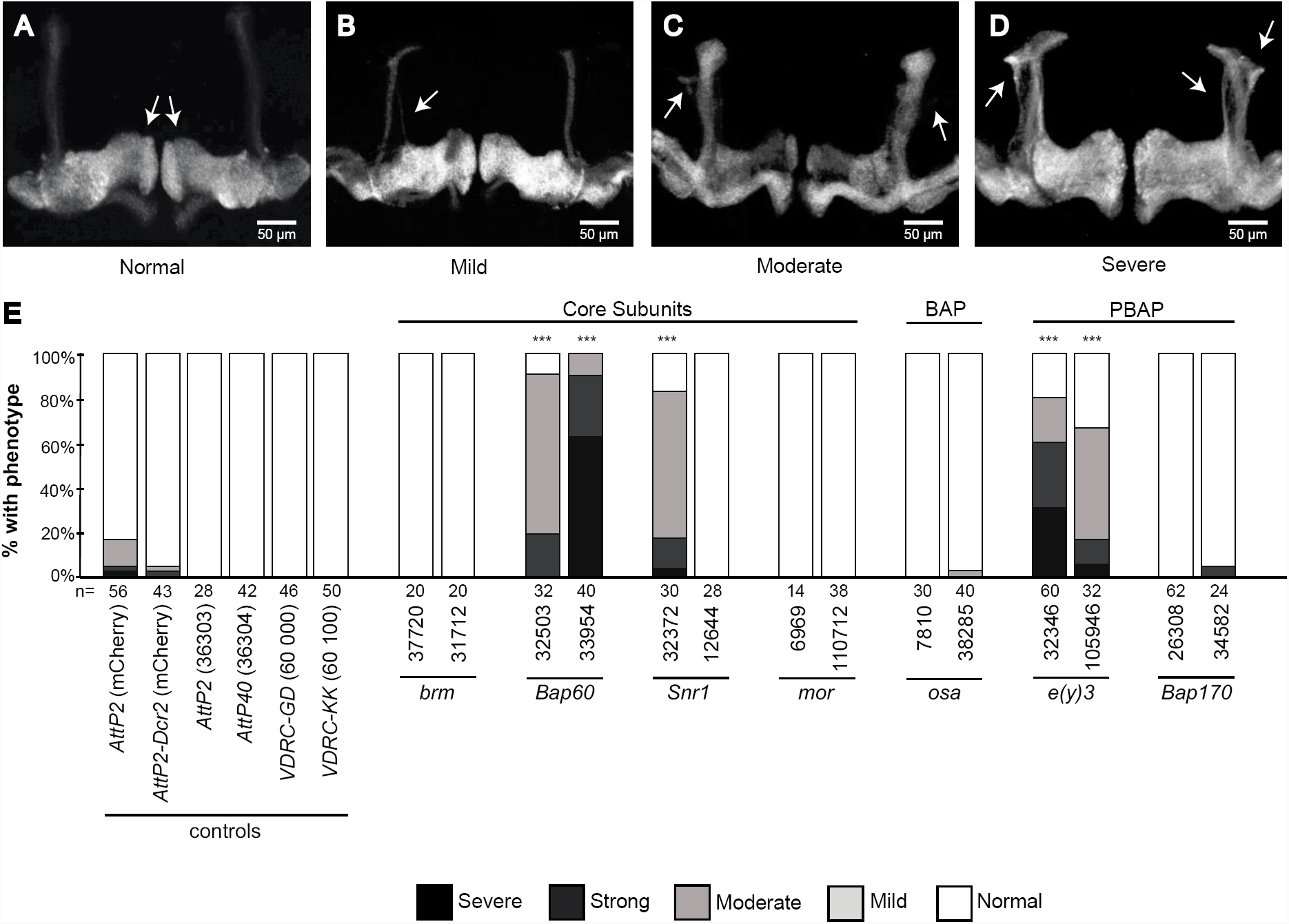
Quantification of extra dorsal projections in SWI/SNF knockdown mushroom bodies. (**A**) The appearance of extra dorsal projections was qualitatively classified into four categories to account for the observed variation in phenotype severity. Confocal projections show representative images for normal MB morphology, as well as the mild, moderate, and severe extra dorsal projection phenotypes. (B) Bar plot showing the total percentage of brains exhibiting normal (white), mild (light gray), moderate (dark gray) and severe (black) extra dorsal projections. The total number of mushroom bodies analyzed for each genotype is indicated below the bar graph. * P<0.05, ** P<0.01, *** P<0.001, Fisher’s exact test, Bonferroni-Dunn test for multiple comparisons.

### The PBAP complex is required for axon pruning during γ neuron remodeling

Previous studies have shown that extra dorsal projections can arise due to defects in γ neuron remodeling that occur during pupal morphogenesis^41-43^. During the larval stages of development, MB γ neurons project both dorsally and medially. During the first 18 hours of pupal development, the dorsal and medial projections are pruned back to the peduncle. This is followed by re-extension of the γ neurons medially, but not dorsally, to form the adult γ lobe^39^ (**Figure 4A**). The appearance of extra dorsal projections in adult flies is often caused by a failure to prune larval γ axons^41-43^. To investigate whether the extra dorsal projections we observed were the result of unpruned γ neurons, and not from aberrantly formed α/β neurons, we performed RNAi knockdown using two γ-neuron specific Gal4 drivers; *MB607B-Gal4* (data not shown) and *MB009B-Gal4* (**Figure 4B**). Indeed, knockdown of Bap60 and e(y)3 using these drivers did result in the appearance of extra dorsal projections in adult flies. Knockdown of Snr1 did not show a strong presence of extra dorsal projections but did show a reduced volume of γ-neurons in general.

**Figure 4:**
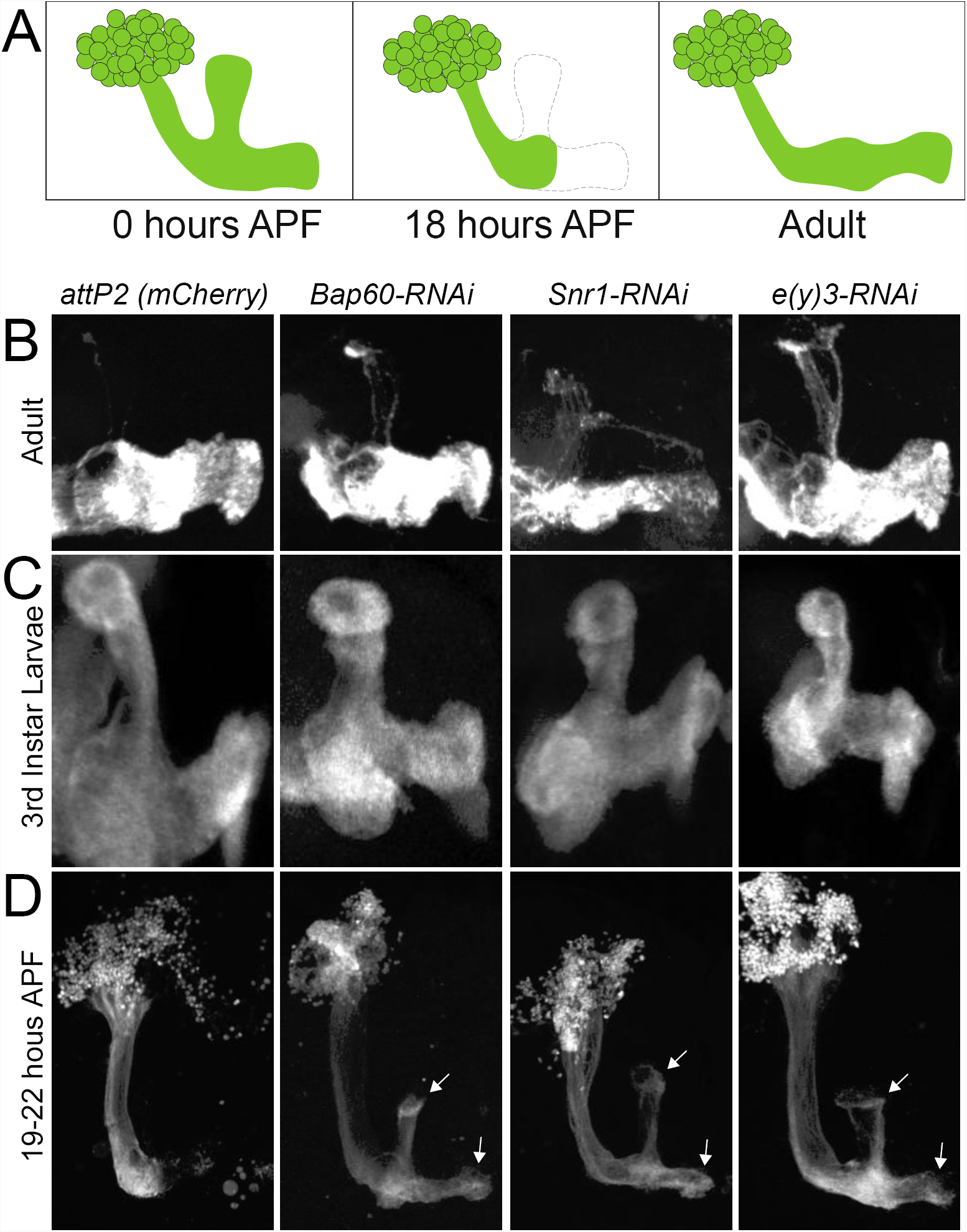
The PBAP complex is required for mushroom body γ-neuron remodeling. (**A**) Schematic diagram of γ-neuron remodeling. (**B-D**) Confocal images showing γ-neuron morphology in controls (expressing an RNAi against mCherry) compared to SWI/SNF knockdown RNai lines for Bap60 Snr1 and e(y)3. Adult γ-neurons were targeted using the γ-specific *MB009B-Gal4* driver. For 3^rd^ instar larvae and pupae, γ-neurons were targeted using *R14H06-Gal4*.

Next, we investigated MB γ lobe structure at the larval and pupal stages to see whether extra dorsal projections resulted from pruning, or aberrant re-extension of γ neurons. Knockdown of Bap60, Snr1, and e(y)3 caused no notable defects in larval MB morphology, suggesting that axon pathfinding can occur normally in knockdown flies as in controls (**Figure 4C**). In pupae, γ neuron pruning was observed at 19-22 hours after pupation in controls, as expected (**Figure 4D**). In contrast, knockdown pupae for Bap60, Snr1, and e(y)3 did not show pruning (**Figure 4D**), verifying that the extra dorsal projection phenotype observed in adult flies is the result of defects in γ neuron pruning during pupal morphogenesis. Interestingly, this phenotype seems to be specifically related to the PBAP complex, since knockdown of the PBAP subunit e(y)3 causes a strong remodeling defect, while knockdown of the BAP specific subunit osa does not (**Figure 3**).

### Faded γ lobes suggest a role for the BAP complex in maintaining survival of mature neurons

The final observed morphological phenotype, described as faded γ lobes, is characterized by normal MB morphology, with a shift in the intensity of fluorescent labeling by GFP. *R14H06-GAL4* is specifically expressed in the α/β and γ neurons of the MB^37^. In controls, GFP expression is strongest within the γ-lobe and weakest within the α/β lobes (**Figure 2A**). For some genotypes, SWI/SNF knockdown caused the appearance of faint γ-lobes that were otherwise morphologically normal. Initial attempts at quantification of this phenotype produced a high level of variability. However, the phenotype was most strongly observed upon RNAi knockdown of brm, and osa. Since our initial screen included flies between 1-7 days of age, we reasoned that the high variability observed in the γ-fade phenotype may be due to the variable age of the flies tested. Therefore, we tested the effects of age in brm and osa knockdown flies. At 1 day, knockdown flies looked similar to controls. However, at 7 days of age γ-lobe fluorescence was clearly and consistently reduced in brm and osa knockdown MBs (**Figure 5A-5C**). This phenotype appears to be specific to the BAP complex as knockdown of the BAP specific subunit osa causes a strong phenotype, while PBAP specific subunits had no effect on γ-lobe intensity.

**Figure 5:**
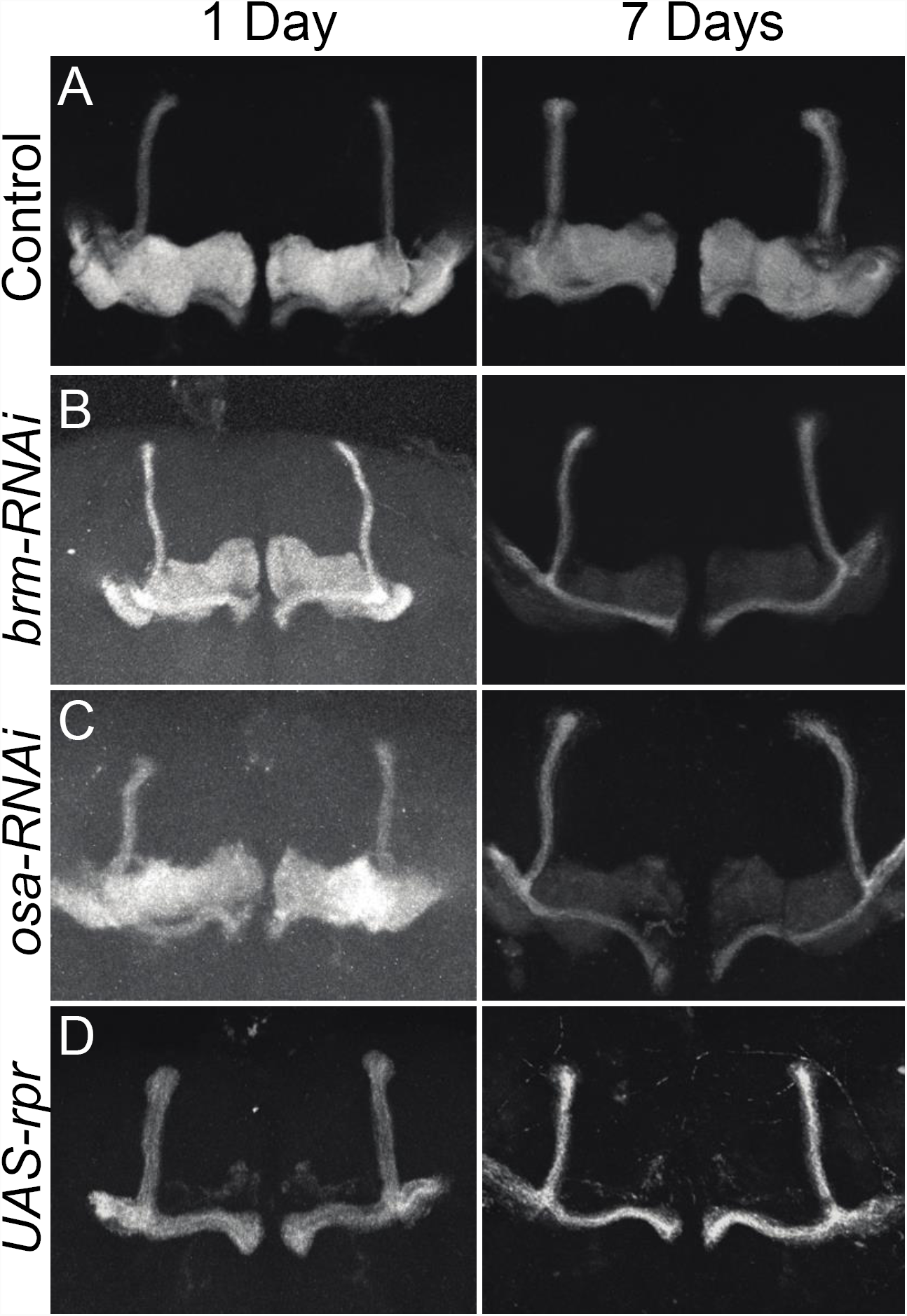
The BAP complex is required for MB γ-neuron survival during ageing. Confocal projections showing MBs labeled with *R14H06-Gal4* and *UAS-mCD8::GFP* at 1 and 7 days after eclosion. Controls expressing and mCherry RNAi (**A**) are compared to flies expressing RNAi constructs targeting brm (*UAS-brm-RNAi*^31712^) (**B**), osa (*UAS-osa-RNAi*^7810^) (**C**), and the cell death gene *UAS-reaper* (**D**).

We reasoned that faded γ lobes may result from a decrease in γ neuron survival due to programmed cell death resulting from SWI/SNF knockdown. To test this, we expressed the cell death gene *reaper* using *R14H04-Gal4. Reaper* expression caused a clear and consistent γ-fade phenotype at 1 and 7 days of age, and similar to SWI/SNF knockdown, did not affect the appearance of the α and β lobes (**Figure 5D**). Although the phenotype was more severe than that seen upon SWI/SNF knockdown, this suggested that increased cell death could underlie the γ-fade phenotype.

### MB-specific SWI/SNF knockdown causes memory defects

Having identified a role for the SWI/SNF complex components in different aspects of mushroom body morphogenesis, we asked whether this might have a functional impact at the organismal level on learning and memory. To test MB-specific SWI/SNF knockdown flies for defects in learning and memory, we used a classic assay called courtship conditioning^44,45^. This assay involves experience based modification of male courtship behaviour towards female flies. Naïve males court females at a high frequency, but pre-mated females are non-receptive and will reject male courtship attempts. After rejection by a non-receptive female, male flies exhibit a learned reduction of courtship behaviour. Short term memory (1 h after rejection) can be induced by a 1 h period of sexual rejection, while long-term memory (24 h after rejection) can be induced by a 7 hour training period. Analysis of short- and long-term courtship memory was performed for six different control genotypes that represent different genetic backgrounds associated with different RNAi lines (see methods). Each of the six control genotypes demonstrated a significant reduction of CI relative to naïve flies for both short- and long-term memory (**Figure S5 and S6**). The average control learning index (LI), which is the proportional reduction in CI due to training, was 0.42 ± 0.0096 for short-term memory, and 0.24 ± 0.017 for long-term memory (**Figure 6**), which is consistent with previous studies^46,47^. As such, courtship conditioning is effective for eliciting both short- and long-term memory across a variety of different control strains that represent different genetic backgrounds of the MB-specific SWI/SNF RNAi knockdown flies.

**Figure 6:**
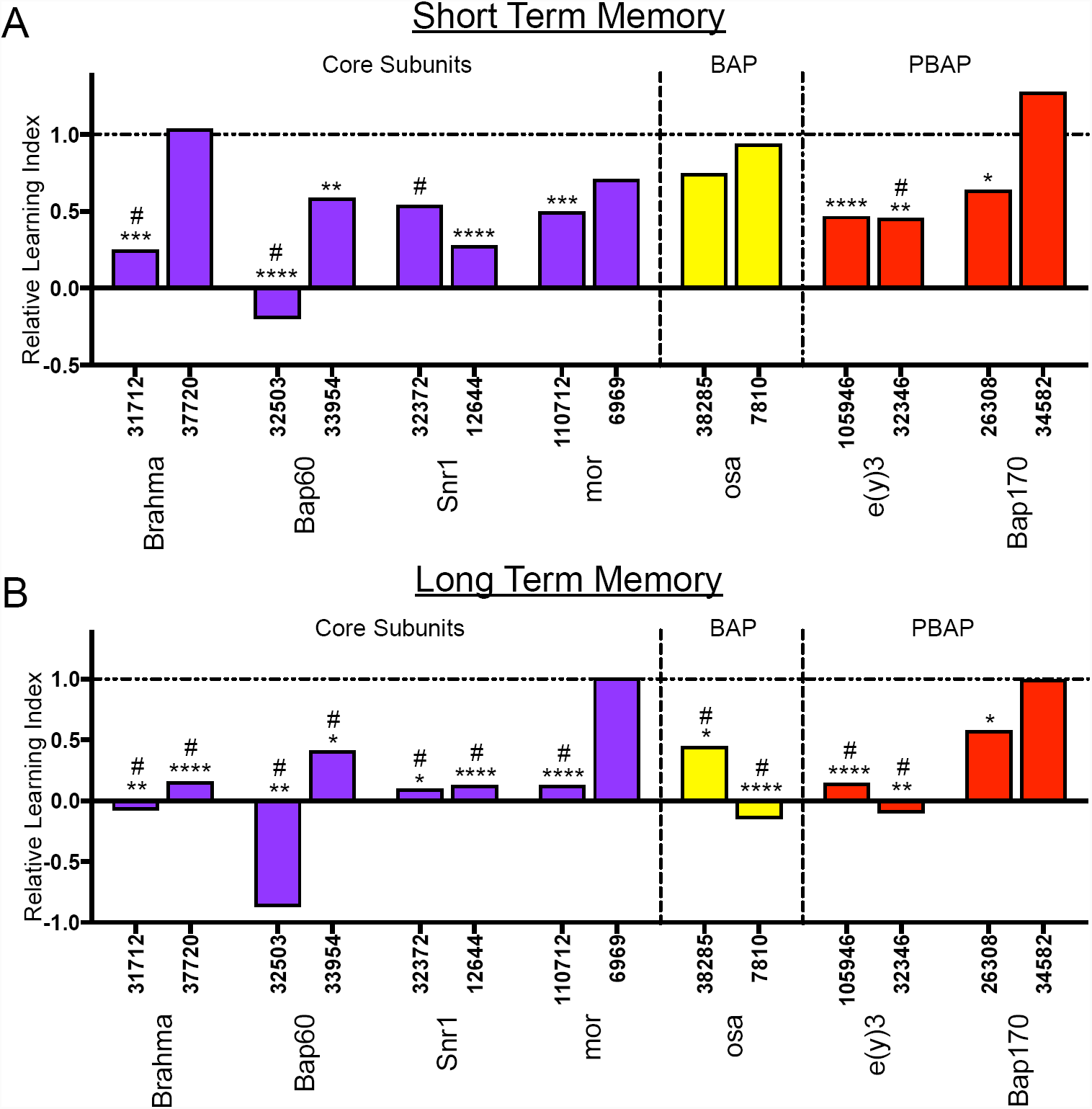
The *Drosophila* SWI/SNF complex is required in the MB for short- and long-term courtship memory. Bar graphs showing the relative learning index (LI) of SWI/SNF RNAi knockdown flies compared to their appropriate genetic background control (see methods) for (**A**) short-term memory and (**B**) long term memory. Blue bars represent Core SWI/SNF subunits, yellow bars represent BAP specific subunits, and red bars represent PBAP specific subunits. # indicates a memory defect indicated by no significant reduction in courtship index (CI) in naïve flies compared to trained flies for that genotype (Kruskal Wallis test, raw CI and LI data available in **Figure S5 and S6**). *p<0.05, **p<0.01, ***p<0.001, ****p<0.0001, randomization test, 10,000 bootstrap replicates.

Next, we examined short and long-term memory in MB-specific SWI/SNF knockdown flies. **Figure 6** shows the relative LI for each RNAi line compared to the appropriate genetic background control (see methods and **Table S2**). The raw courtship data and LIs are shown in **Figures S5 and S6**. Memory defects were widespread among the different SWI/SNF subunits (**Figure 6**). Generally, LTM was more effected, with 86% of RNAi lines inducing LTM loss (**Figure 6B**) compared to 64% for STM (**Figure 6A**). For brm, Bap60, Snr1, osa, and e(y)3, we observed memory defects that were consistent between two RNAi lines. Only mor and Bap170 showed inconsistencies in memory phenotypes between the two RNAi lines, with only one RNAi line inducing memory loss. Together, these data suggest that SWI/SNF expression in the MB is generally important for normal memory.

Memory defects resulting from SWI/SNF knockdown do appear to correlate with morphology defects. Genes that were associated with γ neuron remodeling defects, (*Bap60, Snr1*, and *e(y)3*) showed a consistent loss of STM and LTM (**Figure 6**). Interestingly, while most RNAi lines effect both STM and LTM, only the BAP-specific subunit osa shows an LTM-specific defect. Taken together, these results suggest that PBAP is more broadly required, for STM and LTM, while BAP is specifically required for normal LTM.

## Discussion

Our knowledge of genetic causes of ID is vast. Here, we show that dominant ID genes are highly connected and enriched for functions related to neuronal signaling and chromatin regulation (**Figure 1**). This suggests that many different ID syndromes, which are defined by mutations in different genes, might result from disruption of common cellular and molecular processes. Understanding these common processes represents a significant challenge. Here, we focus on the SWI/SNF complex, which is the most enriched cellular component among dominant ID genes (**Figure 1B**). The role of this complex in cell type specification and differentiation is well understood, however, its function in postmitotic neurons is not extensively investigated^48^. In a screen targeting memory forming neurons of the *Drosophila* MB, we identified a novel role for several individual components of this complex in post-mitotic neuronal processes including, neuron remodeling, survival, and memory (**Figures 3-6**). Interestingly, the PBAP and BAP conformations of the complex seem to have distinct roles in postmitotic neurons. PBAP components are required for morphological remodeling of MBγ neurons during metamorphosis and are involved both short- and long-term memory (**Figures 4 and 6**). In contrast, the BAP complex is required for MBγ neuron survival and has a specific role in long-term memory. This study has revealed new roles for the SWI/SNF proteins in the biology of memory forming neurons and provides a comprehensive phenotypic analysis that will serve as a basis for further investigation into the underlying gene regulatory mechanisms.

### The PBAP complex regulates MBγ neuron remodeling

We show here that the PBAP-specific SWI/SNF subunit e(y)3 and the core subunits Bap60 and Snr1 are required for pruning back of MBγ neurons during the early stages of pupal metamorphosis (**Figure 4**). Interestingly, brm, the ATPase subunit of the core complex has previously been implicated in pruning of multidendrite neurons lining the body wall in the *Drosophila* larvae^49^. Using a dominant negative brm transgene, Kirilly et al. ^49^ also identified a role for brm in MBγ neuron pruning in pupae. We did not observe a MBγ remodeling defect upon brm RNAi knockdown. This suggests that RNAi did not sufficiently reduce brm protein level in order to induce an effect. Indeed, we see that brm RNAi lines reduce mRNA levels to about 50%, suggesting that some protein is likely still produced. The dominant negative brm may be more efficient in silencing SWI/SNF complex activity. Despite this, we did observe clear effects of brm knockdown on memory and neuron survival, suggesting that some processes are more tolerant of reduced brm levels. It is possible that RNAi knockdown of e(y)3, Bap60, and Snr1 results in remodeling defects because these proteins are required for critical protein-protein interactions that are specifically required for remodeling. The remodeling process may be less tolerant to dosage changes in these genes. It is well established that ecdysone signaling is critical for MBγ neuron remodeling^41^. Interestingly, e(y)3 is implicated in direct protein-protein interactions with the ecdysone response protein HR3 in S2 cells^50^, however, whether this mechanism is conserved in the regulation of MBγ neuron pruning remains to be determined.

### A role for the BAP complex in age dependent neuron survival

RNAi knockdown of some SWI/SNF components, including brm and the BAP specific subunit osa, caused an age dependent loss of MBγ neurons (**Figure 5**). This suggests that in some contexts, loss of the SWI/SNF complex may be associated with neurodegeneration or neuronal cell death. There is currently no indication that ID associated SWI/SNF disorders have a neurodegenerative component^20,51,52^, however, this cannot be ruled out as ID is typically diagnosed at a very young age. Interestingly, mutations in the human SWI/SNF gene *SS18L1* have been implicated in amyotrophic lateral sclerosis^53^, a neurodegenerative disorder characterized by loss of motor neurons. Therefore, our analysis targeting postmitotic neurons may have revealed an unappreciated role for the SWI/SNF complex in regulating age dependent neuronal survival. This may be specific for certain types of neurons, as the effect is limited to MBγ neurons and not observed in MBα/β neurons.

### Different roles for BAP and PBAP in memory

Interestingly, we observed that the PBAP specific subunit e(y)3 is required for STM and LTM, while the BAP specific subunit osa, is only required for LTM. This phenotypic divergence highlights the differential function of the two complexes and fits with the different roles that we have observed in regulating neuron remodeling (PBAP) and survival (BAP). But what is the underlying cause of the memory phenotypes? For PBAP, loss of STM and LTM correlates with a strong defect in MBγ remodeling. MBγ remodeling is required for normal short-term courtship memory, but not for long-term, memory^54^. Thus, while PBAP associated STM defects may result from defective MB structure, the underlying cause of LTM defects remains a point of speculation. Both BAP and PBAP could regulate LTM through the regulation of neuronal gene expression effecting the general fitness of neurons. Alternatively, these complexes may be required for neuron activity induced gene expression, which is required for normal LTM, and not STM^55^. Interestingly, the mammalian SWI/SNF complex has recently been shown to mediate enhancer selection in fibroblasts in cooperation with FOS^10^, a critical neuron activity regulated protein involved in memory^56^. This suggests that SWI/SNF components may be essential in enhancer mediated gene activation in response to environmental stimuli, which points to a potential role for the SWI/SNF complex in neuron activity induced gene expression. Our study provides a basis to further investigate this hypothesis in the context of *Drosophila* learning and memory.

## Methods

### Network and GO analysis of dominant ID genes

Network analysis of DIGs was done as described^1^, using annotated interactions from BioGrid 3.2.108 (release January 1, 2014)^34^ and the Human Protein Reference Database^35^ (release 9, April 13, 2010). The PIE score, connectivity and the associated p-values were calculated using the PIE algorithm^36^. GO analysis of DIGs was performed using DAVID, version 6.8^57^. The most enriched GO terms with a Bonferroni corrected p-value <0.05 are shown.

### *Drosophila* Stocks and Crosses

Flies were reared at 70% humidity on a 12h-12h light-dark cycle on standard cornmeal-agar media. All of the RNAi lines and genetic background controls were obtained from either the Transgenic RNAi project (TRiP)^58^ (via the Bloomington *Drosophila* Stock Center), or the Vienna *Drosophila* Resource Center^59^ (listed in **Table S2**). The following additional stocks were obtained from the Bloomington *Drosophila* Stock Center: *UAS-mCD8*::*GFP* (5137), *Pin*^*Yt*^*/CyO, UAS-mCD8::GFP* (5130), *UAS-Dcr-2* (24650), *UAS-reaper* (5824), *R14H06-Gal4* (48667), and *Actin-Gal4/CyO* (25374). The MB γ-lobe specific split-GAL4 lines MB607B and MB009B were obtained from the Fly Light Collection at Janelia Research Campus^60^.

To study the effects of SWI/SNF knockdown in post-mitotic memory forming neurons, the UAS/Gal4 system was used for targeted RNA interference using the mushroom body (MB) specific *R14H06-Gal4* driver line^37^(**Figure 2A**). The RNAi constructs used in this study consist of both short and long hairpin RNA molecules. Dicer-1 is endogenously expressed in the fly, and this expression is effective for the processing of short hairpin RNAs (TRiP’s VALIUM20, VALIUM21 & VALIUM22 collections) ^58,61^. However, long hairpin RNAs require the co-expression of Dicer-2 to achieve optimal knockdown (TRiP’s VALIUM1 & VALIUM10 collections, VDRC’s GD & KK libraries) ^59^. Therefore, *UAS-Dicer-2* was co-expressed in the MB when long hairpin RNA lines were used for knockdown.

For analysis of MB morphology, RNAi lines (**Table S2**) were crossed to flies of the genotypes *UAS-mCD8::GFP/CyO; R14H06-GAL4/TM6* or *UAS-Dcr2/CyO; R14H06-GAL4, UAS-mCD8::GFP/TM6*, at 29°C. For courtship conditioning experiments, RNAi lines (**Table S2**) were crossed to *R14H06-Gal4* or *UAS-Dcr2/CyO; R14H06-GAL4/TM6*, at 25°C. For all knockdown experiments, control flies were generated by crossing the appropriate genetic background strain to the driver line (see **Table S2** for controls used for each RNAi line). For knockdown experiments using TRiP RNAi lines, the corresponding controls were generated using either *attP2(36303), attP40(36304)*, or *attP2(mCherry)*. For knockdown experiments using RNAi lines obtained from the VDRC, controls were generated using *VDRC-GD(60000)* or *VDRC-KK(60100)*.

### Validation of RNAi lines by qPCR and lethality assay

As a simple phenotypic test to assess RNAi efficiency, we measured survival upon ubiquitous knockdown, with the knowledge that null mutations in most genes investigated in this study cause lethality. *Actin-GAL4/CyO* flies were crossed to *UAS-RNAi* flies for *brm, Bap60, Snr1, mor, Bap111, osa, e(y)3, Bap55, polybromo,* and *Bap170* (**Table S1**). Percent survival was calculated by comparing the number of progeny with straight wings to the number of flies with curly wings (not receiving *Actin-Gal4*). Validation of RNAi knockdown efficiency by qPCR was done as described^38^, using polyA purified RNA as a template and primers directed towards the 5’ side of the predicted RNAi induced cleavage site. Western blotting was performed according to standard protocols using Bap111^33^ (1:2000) and a mouse anti-β-tubulin (1:8000; Developmental Studies Hybridoma Bank) primary antibodies with goat anti-guinea pig (1:8000) and goat anti-mouse (1:3000) horseradish peroxidase (HRP)-conjugated secondary antibodies.

### Analysis of MB morphology

Adult brains expressing *UAS-mCD8::GFP* with *R14H06-Gal4* were dissected in PBS and fixed with 4% paraformaldehyde for 45 minutes at room temperature before mounting in Vectashield (Vector Laboratories) and imaging using a Zeiss LSM 510 duo vario confocal microscope. Confocal stacks were generated using 100 µm slices and processed using Image J software (Fiji)^62^ and Adobe Photoshop.

Gross MB morphology was assessed and qualitatively quantified by examining confocal stacks. Five distinct morphological variations were observed, including 1) missing α and β lobes, 2) β-lobe fibers crossing the midline, 3) extra dorsal projections, 4) stunted γ-lobes, and 5) faded γ-lobes. For missing lobes, each brain was scored as either missing a lobe or not. Due to variability in the severity, β-lobe crossing, extra dorsal projections, and stunted γ-lobes were classified as ‘normal’, ‘mild’, ‘moderate’, ‘strong’ or ‘severe’ as indicated in **Figure S3**, **Figure S4**, and **Figure 3**. The β-lobe crossing phenotype was scored based on the width and density of GFP labelled β-lobe fibers crossing the midline. Extra dorsal projections were classified based on the number and thickness of dorsal projections adjacent to the α-lobe. Stunted γ-lobes were classified based on the relative size of the γ-lobe.

### Courtship conditioning

Courtship conditioning was performed as described previously^44^. Male knockdown flies were collected at eclosion and raised in isolation for five days before pairing with a pre-mated female for one hour, for short-term memory experiments, or seven hours, for long-term memory experiments. During training, PMFs reject male courtship attempts. Following training, males were placed in isolation for one hour, for short-term memory experiments, or 24 hours, for long-term memory experiments. Following isolation, both naïve and trained males were individually paired with a new premated female in 18 well courtship chambers and courtship behaviour was recorded for 10 minutes. Experiments were conducted on at least three different days with a maximum of 18 male female pairs per condition per day. For each of the males used in this study a Courtship Index (CI) was calculated through manual scoring of courtship behaviour by expert observers.

Statistically, loss of memory was identified using two complimentary methods. Reduction of the mean CI of trained (CI_trained_) flies compared to naïve (CI_naive_) of the same genotype was compared using a Kruskal-Wallis test followed by pairwise comparisons using Dunn’s test. No significant reduction in the mean CI due to training (p>0.05) indicates a defect in memory. In some cases, a group of flies may show a significant reduction in CI due to training, but still show less capacity for memory compared to controls. For this comparison a learning index (LI) was calculated (LI=(CI_naive_-CI_trained_)/CI_naive_). LIs were compared between genotypes using a randomization test^63^ (10,000 bootstrap replicates) using a custom R script^44^ and the resulting p-values were corrected for multiple testing using the method of Bonferroni.

## Acknowledgments

We thank the Bloomington Drosophila Stock Center, the Vienna Drosophila Resource Center, and the Transgenic RNAi Project at Harvard Medical School for generating and distributing fly stocks used in this study. Thanks to M. Fenckova and P. Cizek for providing the protein interaction enrichment (PIE) analysis program. The Bap111 antibody was a gift from C.P. Verrijzer. This work was funded by the National Science and Engineering Research Council of Canada, the Canadian Institute of Health Research, the Canada Research Chairs program, and the Canadian Foundation for Innovation.

---

**Figure S1:**
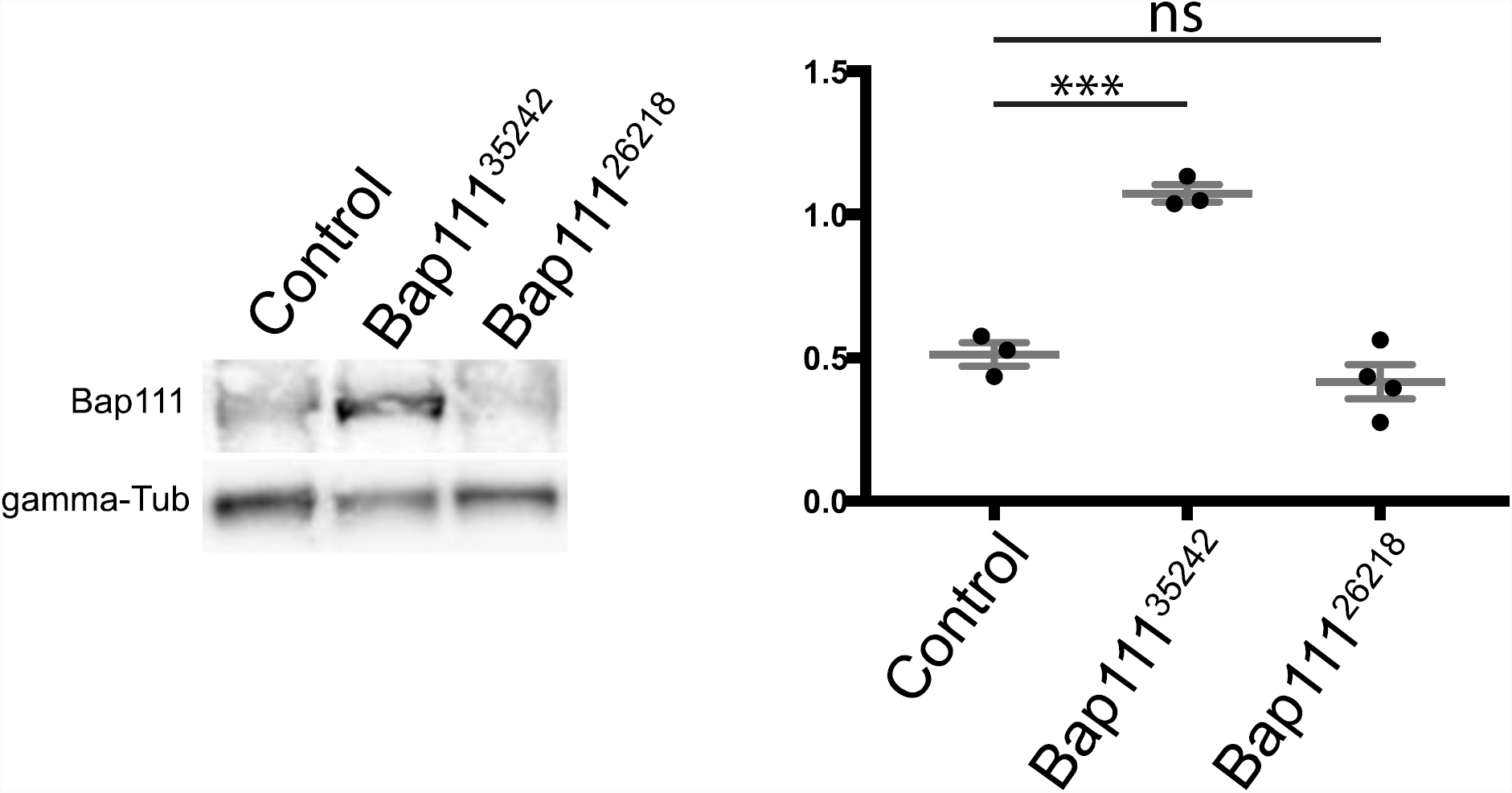
Quantification of Bap111 expression in RNAi knockdown larvae by Western blot. Representative bands for Bap111 and the loading control gamma-tubulin (left). At least three biological replicates were quantified using image J (right). *** p < 0.001 - ANOVA – Dunnett’s test for multiple comparisons.

**Figure S2:**
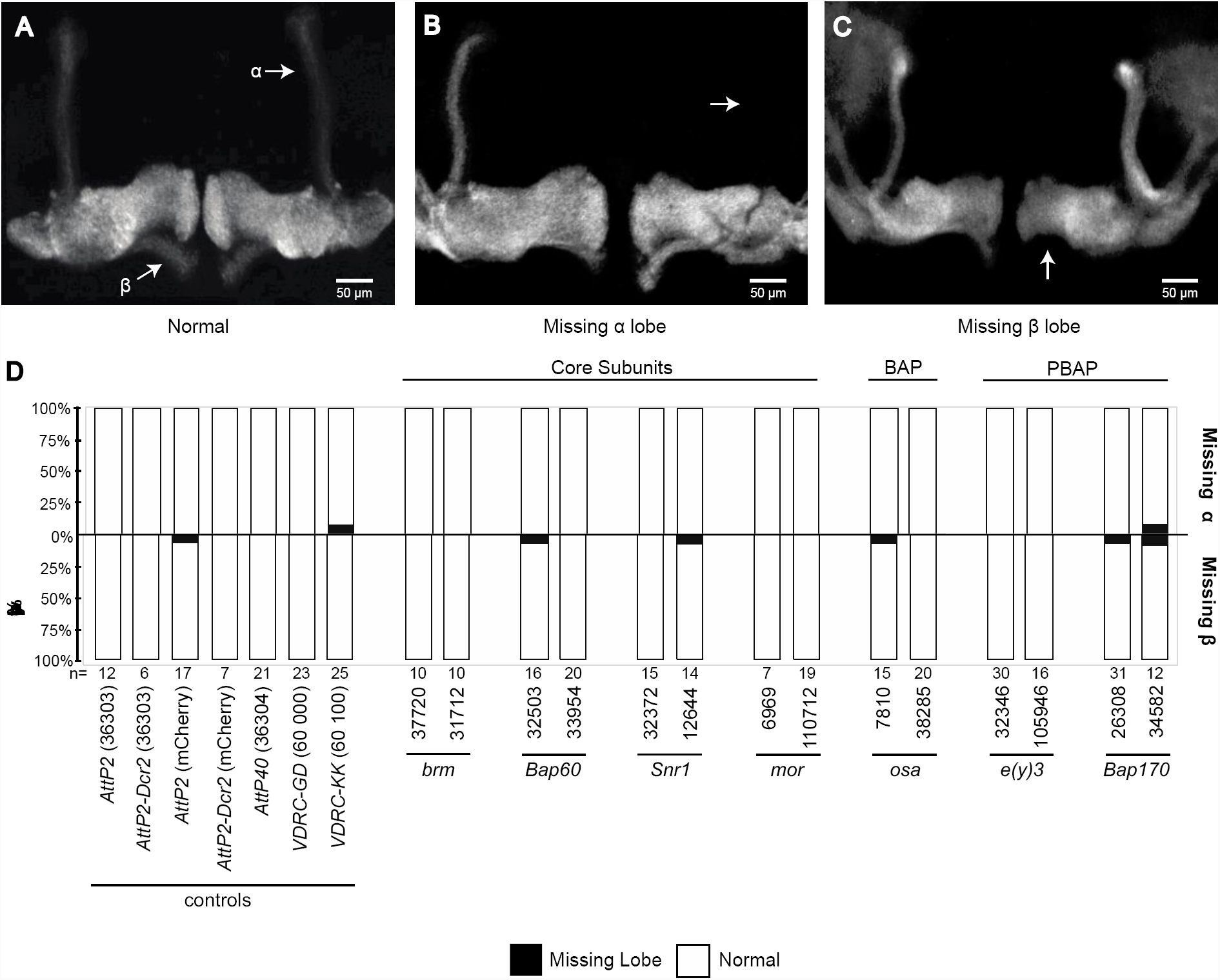
Missing α and β lobes were observed at a low penetrance in SWI/SNF knockdown MBs and controls. Confocal projections show **(A)** normal MB morphology, **(B)** the missing α-lobe phenotype and **(C)** the missing β-lobe phenotype. **(D)** Bar plots indicate the total percentage of brains exhibiting the missing α-lobe phenotype (above x-axis) and missing β-lobe phenotype (below x-axis). The total number of flies analyzed for each genotype is indicated.

**Figure S3:**
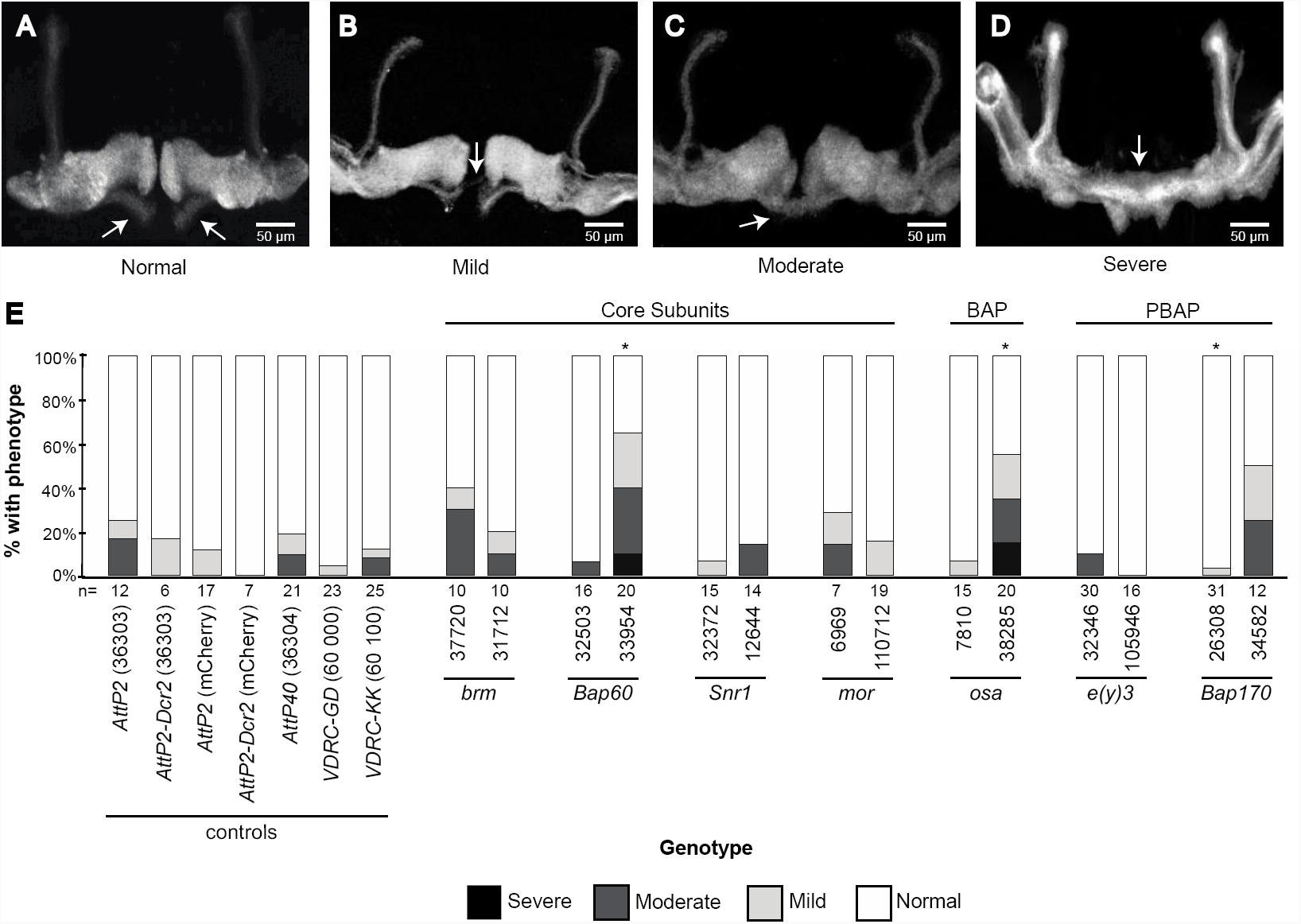
The appearance of β-lobe fibers crossing the midline was observed at a variable penetrance in both knockdowns and controls. The β-lobe crossing phenotype was qualitatively classified into four categories to account for the observed variation in phenotype severity. Confocal projections show **(A)** normal MB morphology, in addition to the **(B)** mild, **(C)** moderate, and **(D)** severe forms of the β-lobe crossing phenotype. **(E)** Bar plot shows the total percentage of brains exhibiting normal MB morphology (white), in addition to the mild (light gray), moderate (dark gray) and severe (black) forms of the β-lobe crossing phenotype. The total number of flies analyzed for each genotype is indicated. The Fisher’s exact test (two tailed) was used to compare the proportion of MBs exhibiting abnormal morphology (sum of mild, moderate and severe proportions) to the proportion exhibiting normal morphology between each knockdown and the appropriate control. * p< 0.05 - Bonferroni-Dunn test for multiple comparisons.

**Figure S4:**
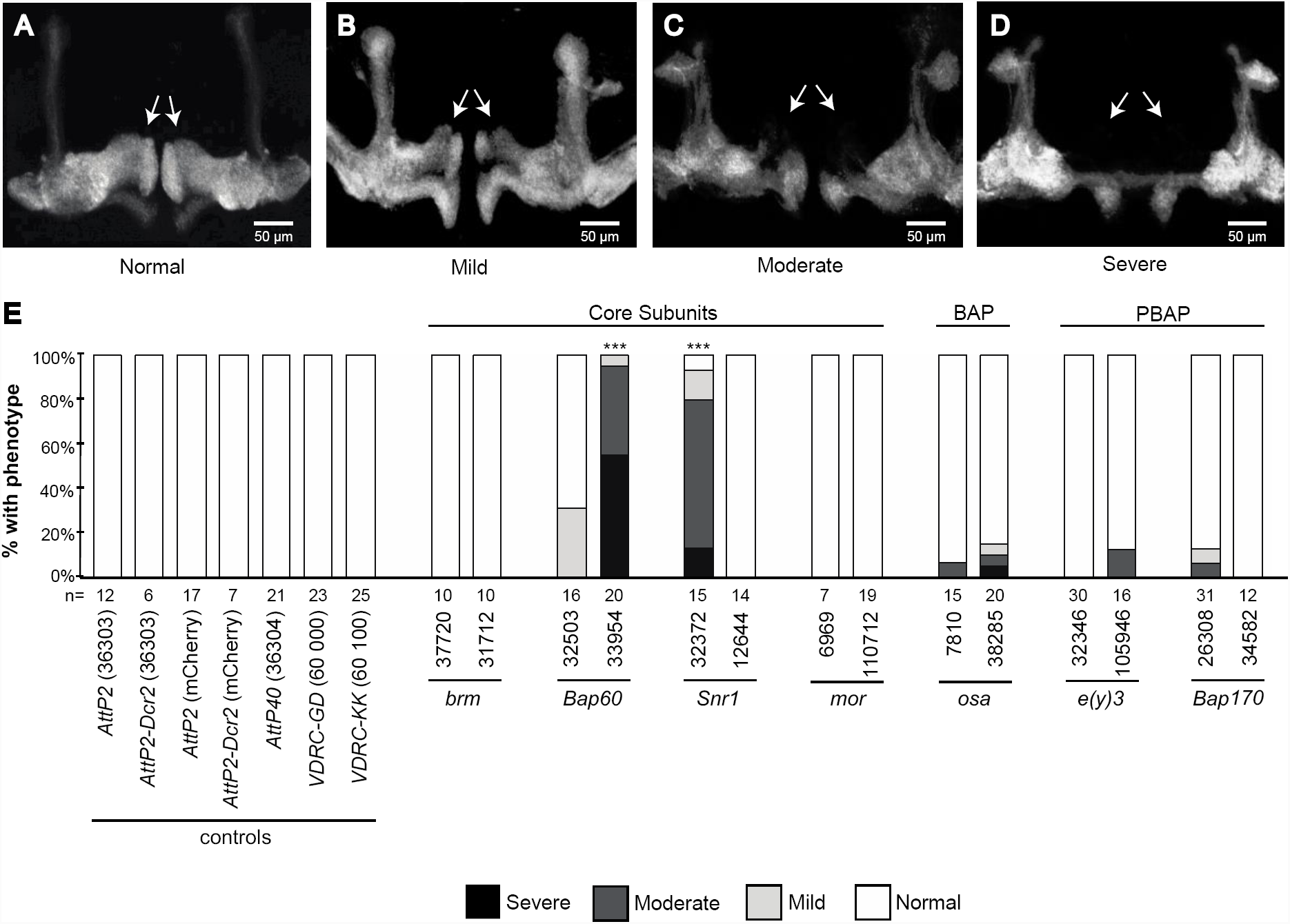
Quantification of stunted γ-lobe in SWI/SNF knockdown mushroom bodies. (**A**) The appearance of stunted γ-lobe was qualitatively classified into four categories to account for the observed variation in phenotype severity. Confocal projections show representative images for normal MB morphology, as well as the mild, moderate, and severe stunted γ lobe. (**B**) Bar plot showing the total percentage of brains exhibiting normal (white), mild (light gray), moderate (dark gray) and severe (black) phenotypes. The total number of mushroom bodies analyzed for each genotype is indicated. *** P<0.001, Fisher’s exact test, Bonferroni-Dunn test for multiple comparisons.

**Figure S5:**
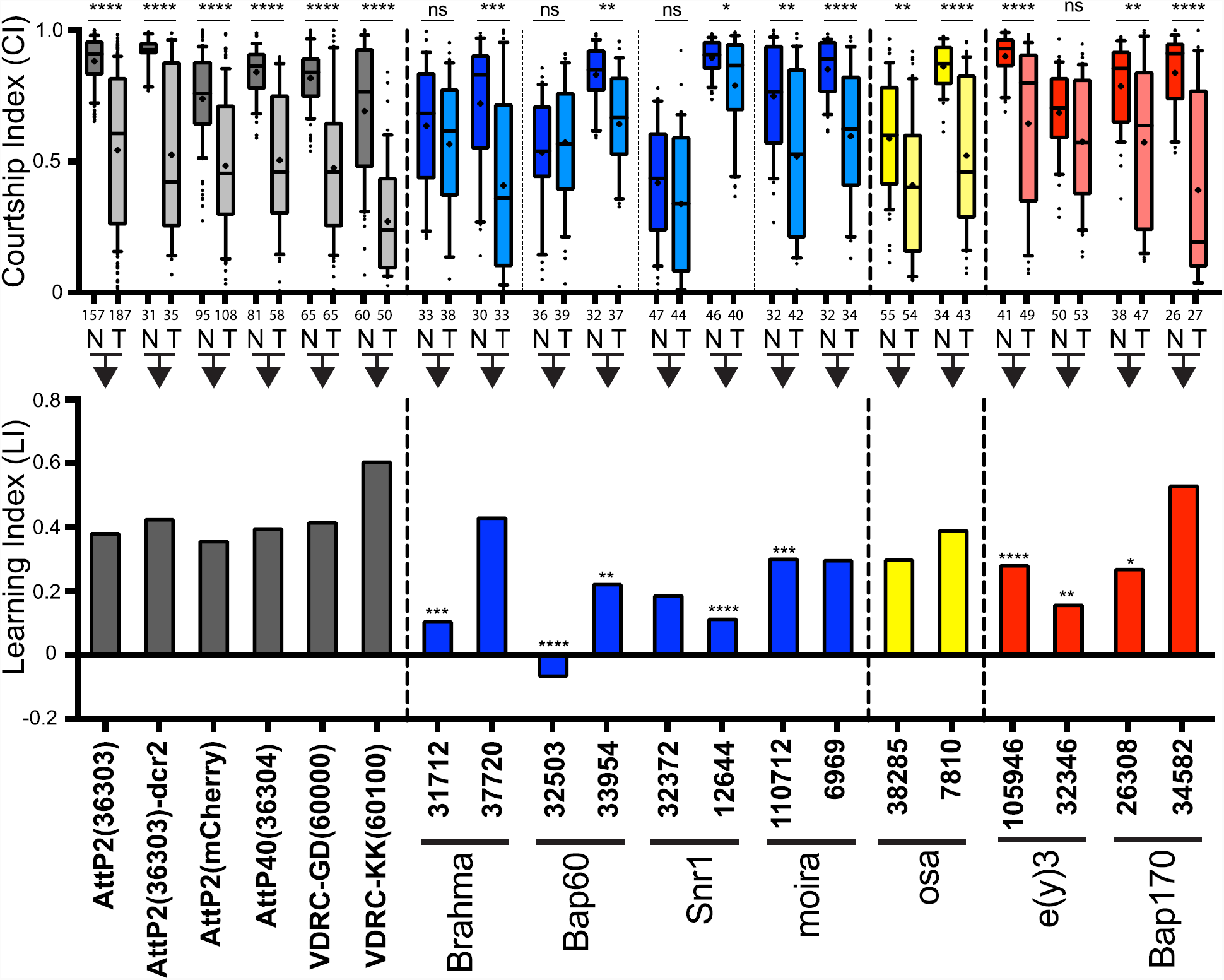
MB-specific SWI/SNF knockdown causes defects in short-term courtship memory. (**A**) Box plots indicating the courtship index (CI) for naïve (N) flies and flies that were trained for short-term courtship memory by exposure to sexual rejection for 1 hour (T). Trained and naive flies were tested at the same time, 1 hour after rejection. *p<0.05, **p<0.01, ***p<0.001, ****p<0.0001, Kruskal Wallis and Dunn’s test. (**B**) Bar graphs showing the learning index (LI) for each genotype (LI = (CI_naive_- CI_trained_)/CI_naive_). *p<0.05, **p<0.01, ***p<0.001, ****p<0.0001, randomization test, 10,000 bootstrap replicates. Grey bars - controls, blue bars-SWI/SNF core subunits, yellow bars – BAP specific, red bars – PBAP specific. Contol genotypes, SWI/SNF genes, and RNAi stock numbers are indicated.

**Figure S6:**
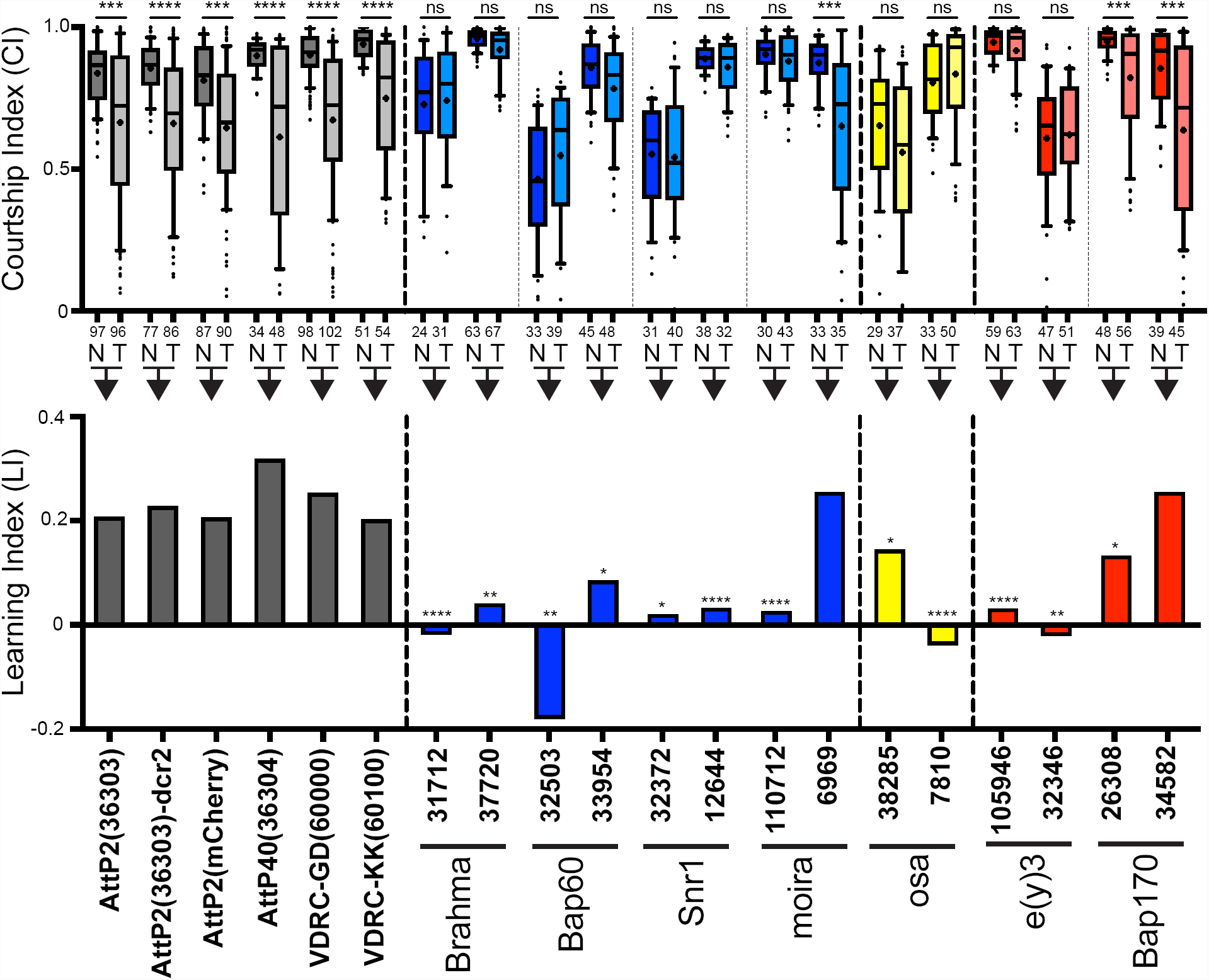
MB-specific SWI/SNF knockdown causes defects in long-term courtship memory. (**A**) Box plots indicating the courtship index (CI) for naïve (N) flies and flies that were trained for short-term courtship memory by exposure to sexual rejection for 7 hours (T). Trained and naive flies were tested at the same time, 24 hour after rejection. *p<0.05, **p<0.01, ***p<0.001, ****p<0.0001, Kruskal Wallis and Dunn’s test. (**B**) Bar graphs showing the learning index (LI) for each genotype (LI = (CI_naive_-CI_trained_)/CI_naive_). *p<0.05, **p<0.01, ***p<0.001, ****p<0.0001, randomization test, 10,000 bootstrap replicates. Grey bars - controls, blue bars-SWI/SNF core subunits, yellow bars – BAP specific, red bars – PBAP specific. Contol genotypes, SWI/SNF genes, and RNAi stock numbers are indicated.

**Table S1:** Evaluation of transgenic RNAi lines using a lethality assay and qPCR.

| Gene | RNAi Stock # | Survival Assay |  | mRNA |  |
| --- | --- | --- | --- | --- | --- |
|  |  | % Survival<sup>1</sup> | p-value<sup>2</sup> | qPCR<sup>1</sup> | p-value<sup>3</sup> |
| *mCherry* | 35785 | 80.2 ± 8.40 | < 0.0001 |  |  |
| *Bap170* | 26308 | 1.70 ± 3.70 | < 0.0001 | 45 ± 3.3 | < 0.0001 |
|  | 34582 | 0.71 ± 1.15 | < 0.0001 | 44 ± 9.3 | < 0.0001 |
| *Bap55* | 24703 | 5.00 ± 4.40 | < 0.0001 | NA | NA |
|  | 31708 | 83.3 ± 23.3 | 0.99 | NA | NA |
| *Bap60* | 32503 | 0 | < 0.0001 | 48 ± 13 | 0.0003 |
|  | 33954 | 17.8 ± 12.4 | < 0.0001 | xx | - |
| *brm* | 31712 | 0 | < 0.0001 | 46 ± 4.0 | < 0.0001 |
|  | 37720 | 0.96 ± 0.79 | < 0.0001 | 60 ± 3.0 | 0.0082 |
|  | 34520 | 77.8 ± 15.6 | 0.99 | 97 ± 11 | 0.99 |
| *Bap111* | 35242 | 3.55 ± 0.97 | < 0.0001 | 284 ± 13 | < 0.0001 |
|  | 26218 | 1.41 ± 1.96 | < 0.0001 | 85 ± 8.7 | 0.98 |
| *e(y)3* | 105946 | 0 | < 0.0001 | 35 ± 3.3 | < 0.0001 |
|  | 32346 | 0 | < 0.0001 | 15 ± 3.2 | < 0.0001 |
| *mor* | 110712 | 1.20 ± 1.33 | < 0.0001 | 28 ± 2.1 | < 0.0001 |
|  | 6969 | 0.83 ± 0.93 | < 0.0001 | 22 ± 0.8 | < 0.0001 |
| *Snr1* | 32372 | 4.04 ± 3.92 | < 0.0001 | xx | - |
|  | 12644 | 53.8 ± 18.8 | 0.0040 | 62 ± 3.7 | 0.017 |
| *osa* | 38285 | 2.47 ± 1.55 | < 0.0001 | 59 ± 8.0 | 0.023 |
|  | 7810 | 2.13 ± 0.98 | < 0.0001 | xx | - |

<sup>1</sup>± standard error of the mean
<sup>2</sup>ANOVA with Dunnett test for multiple comparisons
<sup>3</sup>ANOVA with Tukey test for multiple comparisons

**Table S2.**
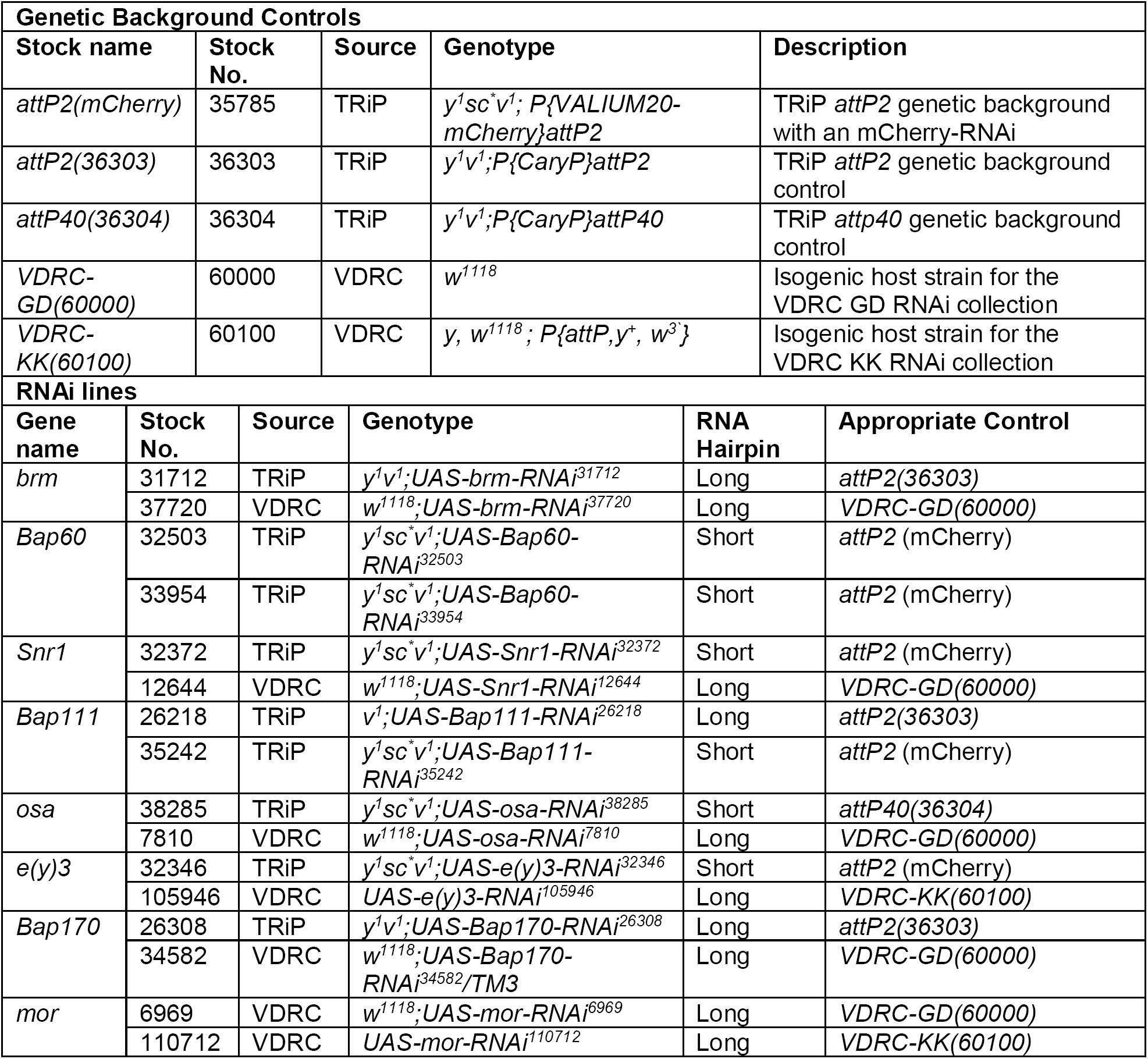
RNAi stocks and genetic background controls used in this study

